# Coordination of pupil and saccade responses by the superior colliculus

**DOI:** 10.1101/2020.08.20.247668

**Authors:** Chin-An Wang, Douglas P. Munoz

## Abstract

The appearance of a salient stimulus evokes saccadic eye movements and pupil dilation as part of the orienting response. Although the role of the superior colliculus (SC) in saccade and pupil dilation has been established separately, whether and how these responses are coordinated remains unknown. The SC also receives global luminance signals from the retina, but whether global luminance modulates saccade and pupil responses coordinated by the SC remains unknown. Here, we used microstimulation to causally determine how the SC coordinates saccade and pupil responses, and whether global luminance modulates these responses by varying stimulation frequency and global luminance in male monkeys. Stimulation frequency modulated saccade and pupil responses, with trial-by-trial correlations between the two responses. Global luminance only modulated pupil, but not, saccade responses. Our results demonstrate an integrated role of the SC on coordinating saccade and pupil responses, characterizing luminance independent modulation in the SC, together elucidating the differentiated pathways underlying this behavior.

## Introduction

A series of responses, including saccadic eye movements and pupil dilation, are evoked by the presentation of a salient stimulus as part of the orienting response ^1–3^. Recent evidence, particularly in monkeys, has shown that pupil dilation is evoked by presentation of a visual target, enhanced by acoustic stimuli, modulated by stimulus saliency, and triggered by microstimulation of the superior colliculus (SC) ^4–7^, a midbrain sensorimotor center that is phylogenetically conserved and causally involved in producing orienting responses ^8–12^. Pupil size and saccadic eye movements are becoming popular and promising indices of cognitive and disease processes ^13^. Are the neural mechanisms underlying the cognitive modulation for pupil size similar to those for saccades? Further, what is the role of the SC in coordinating saccade and pupil responses?

The SC is organized into a retinotopic map of contralateral visual space and has anatomically and functionally differentiated superficial (SCs) and intermediate/deep (SCi) layers ^14^. The SCs receives visual signals from the retina and visual cortex, whereas the SCi receives multisensory and cognitive inputs from various cortical and subcortical areas ^15–18^, including structures and neuromodulatory systems involved in the control of pupil size, such as the locus coeruleus (LC), frontal eye field (FEF), and cholinergic projections from the pedunculopontine tegmental nucleus (PPTN) ^19–23^. Moreover, the SCi projects to the premotor circuitry in the brainstem that initiates the orienting response ^8,10,24^. Electrical microstimulation of the SCi in monkeys induces saccades ^25^, and the properties of the evoked movements are modulated by stimulation parameters and the location of stimulation within the SC, with larger saccade responses (e.g., peak velocity) evoked using stronger stimulation parameters and larger amplitude saccades evoked by moving toward the caudal SCi ^26^. Weak microstimulation in the SCi can evoke pupil dilation without inducing saccades ^4,22^. If the saccade and pupil responses are coordinated by the SCi (referred to as the common drive hypothesis), then varying stimulation parameters and changing stimulation sites should modulate not only saccades, but also pupil responses.

Besides orienting, pupil size also changes to regulate the amount of light entering the retina to optimize visual sensitivity and acuity ^27,28^, with smaller pupil sizes for higher luminance levels, and larger pupil size for lower luminance levels. This luminance response is controlled by the balance of activity between the parasympathetic and sympathetic nervous systems ^29,30^. Specifically, higher parasympathetic activity occurs during higher luminance levels causing pupil constriction, while higher sympathetic activity occurs during lower luminance levels leading to pupil dilation. The SCs receives direct retinal projections including those from intrinsically photosensitive retinal ganglion cells (ipRGCs) ^31^, which are important for luminance encoding and regulating the pupil light reflex ^32–35^. Although visual signals such as visual contrast play an important role in the SC ^36^, the functional role of global luminance signals in the SC is yet to be explored.

Using microstimulation of the SCi with a systematic variation of stimulation frequency and background luminance, we investigate the role of the SCi in coordinating saccade and pupil responses at different luminance levels. If the SCi coordinates saccades and pupil responses, then responses evoked by SCi microstimulation should be correlated on trial-by-trial basis. Additionally, if the SCi is modulated by luminance levels, then background luminance should affect both pupil and saccade responses evoked by SCi microstimulation (Fig 1A). If, by contrast, the luminance signals sent to the SCi are functionally insignificant, then only the evoked pupil responses will be modulated ^29^.

**Figure 1.**
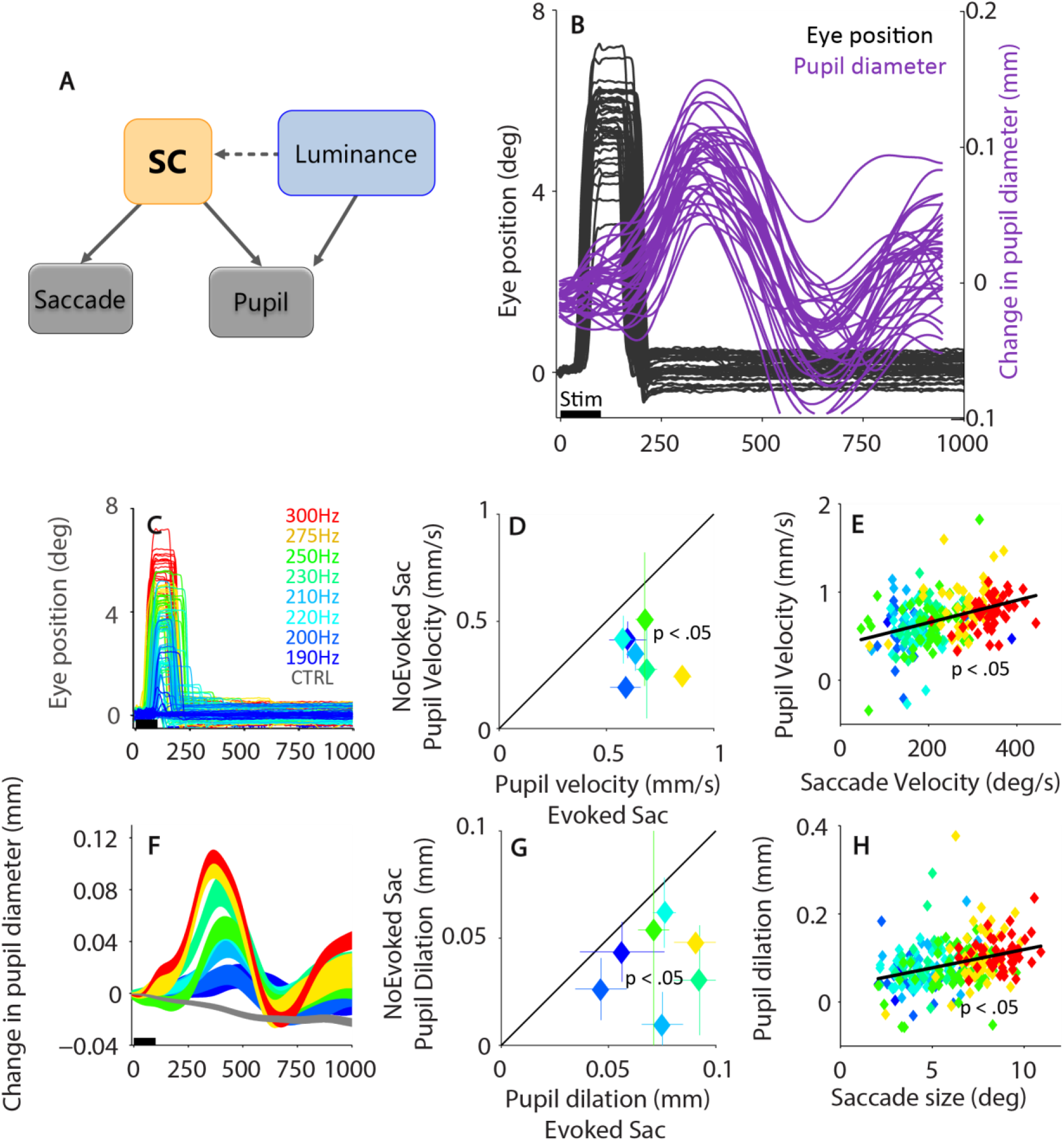
Effect of SCi microstimulation on saccade and pupil responses from an example site. (A) Illustration of hypotheses on the modulation of the superior colliculus on saccade and pupil responses by luminance. (B) Temporal dissociation between evoked saccades (only horizontal eye position) and pupil responses. (C,F) Effects of stimulation frequency on (C) saccade and (F) pupil responses. (D,G) Pupil responses between trials with evoked saccades and trials without evoked saccades in (D) velocity and (G) magnitude. (E,H) Trial-by-trial correlation between normalized pupil and saccade responses in (E) velocity and (H) magnitude. In C,F, the black bar on X-axis indicates the time line of microstimulation. The shaded colored regions surrounding the pupil response represent ± standard error range in different conditions. CTRL: control (no stimulation). Stim: microstimulation.

## Results

### Correlation between saccade and pupil responses via varying train frequency

If the mechanisms of initiation of saccade and pupil dilation are shared through the SCi, then the comparable modulation of saccade and pupil responses by varying stimulation frequency should be observed. More specifically, the common-drive hypothesis postulates that the same efferent neurons in the SCi project to both saccade and pupil control circuits. Therefore, trial-by-trial correlations between saccade and pupil responses induced by SCi microstimulation should be observed. For example: if there are stronger output signals from the SCi, this signal should drive larger saccade and pupil responses. Moreover, because of a gating role of the omnipause neurons e.g., ^37^, signals must surpass the threshold to evoke saccades. Thus, pupil dilation on trials with an evoked saccade should be larger than that on trials without an evoked saccade. To test these predictions, monkeys were trained to perform a simple fixation task in which they were required to maintain fixation upon a central fixation point (FP). Microstimulation was delivered to 24 sites in the intermediate layers (SCi) (9 in monkey A, 15 in monkey B) on 50% of trials (see Materials and Methods for details). Figure 1B shows an example of individual saccade and pupil responses induced by SCi microstimulation (300 Hz, 40 µA, 100 ms). Microstimulation evoked saccades (black traces in Fig. 1B) towards the location that corresponds to the stimulated SC position with a mean latency of 42 ms (22 - 63 ms) after stimulation onset. The monkey then quickly returned his eyes to the central FP with a mean latency of 125 ms (87 - 152 ms) measured after the previous saccade offset. Pupil dilation was also evoked after microstimulation, but the pupil response onset latency was slower, starting near 200 ms and peaking near 300 ms after the stimulation, due to the longer efferent delay of the pupil control circuit ^4^. We focused here on initial pupil dilation because pupil dilation has long been characterized as a component of orienting ^1,38^. Moreover, pupil dilation dynamics evoked by SCi microstimulation ^4^ are similar to those evoked by salient stimuli ^6^, suggesting that similar orienting processes may be involved between two observed pupil dilation. Also, pupil dilation is related to aspects of cognition ^39,40^. Eye movements and variations in eye position could diminish the accuracy of pupil size measurements. Because the eyes of monkeys almost always returned to central FP within 300 ms after stimulation onset (Fig. 1B), to maintain the accuracy of pupil measurement, we measured pupil peak velocity and dilation (which occurred at around 300 to 500 ms after stimulation onset) to correlate pupil responses with saccade responses (see Materials and Methods).

Changes in stimulation frequency altered saccade and pupil responses in a systematic and coordinated manner, as shown in Fig. 1C-H for this example stimulation site. Consistent with the literature ^26^, increasing stimulation frequency systematically increased the probability of evoking a saccade (low to high frequency: 13, 33, 48, 64, 81, 92, 98, 100 %), decreased the latency (low to high frequency, mean value: 86.3, 72.1, 69.3, 78.2, 67.2, 54.0, 41.7, 41.8 ms), and increased saccade amplitude (Fig. 1C, low to high frequency conditions: 3.8, 4.1, 4.6, 4.2, 4.7, 6.1, 7.6, 8.7 deg). Microstimulation also resulted in transient pupil dilation, followed by constriction (purple traces in Fig. 1B). Importantly, these pupil responses were also systematically modulated by stimulation frequency, with larger peak dilation observed for higher stimulation frequencies (Fig. 1F, low to high frequency in magnitude: 0.048, 0.052, 0.080, 0.073, 0.099, 0.072, 0.105, 0.109 mm).

To examine the trial-by-trial correlation between saccade and pupil responses, we divided pupil responses according to the likelihood of evoking saccades. Trials with saccades were associated with larger pupil velocities and greater dilation (Fig. 1D velocity, *t*(6) = 5.05, *p* < 0.01; Fig. 1G magnitude, *t*(6) = 3.86, *p* < 0.01). We also examined the correlation between saccade and pupil responses across individual trials when a saccade was evoked and found that trials with larger saccade responses had larger pupil responses (Fig. 1E in velocity, R = 0.4, *p* < 0.001; Fig. 1H in magnitude, R = 0.34, *p* < 0.001).

Saccade and pupil responses evoked by microstimulation were highly correlated across our sample of stimulation sites (N=24) in two monkeys (Fig. 2). Stimulation frequency systematically modulated the probability of evoking a saccade (Fig. 2A). A generalized linear model was used to characterize this relationship by applying a binomial distribution (saccade evoked or not: 1 or 0) and a logistic link function (a logistic fit). The fitted model yielded an R-squared value of 0.99 (data from data source Fig. 2), suggesting that saccade probability can almost be fully explained by the model of using stimulation frequency as a predictor. Fig. 2B summarizes pupil dynamics following microstimulation (22 sites for which we had four or more stimulation frequency conditions were included in Fig. 2B, 2 sites that only had three stimulation frequency conditions were thus excluded from analysis, N= 22; individual monkey data provided in Supplementary Figure 1, revealing the same stimulation frequency modulation in two monkeys), showing that pupil dilation scaled with stimulation frequency (high to low frequency and control: mean pupil size at an epoch of 250 to 350 ms after stimulation onset: 0.075, 0.053, 0.041, 0.021, −0.006 mm). To illustrate the results across microstimulation sites, we normalized the data by dividing saccade/pupil values by the median value in the highest frequency condition (see Materials and Methods). The common drive hypothesis (Fig. 1A) predicts that pupil responses should be larger on trials in which saccades were evoked, and this is what we observed. Pupil responses evoked by microstimulation were larger on trials with saccades, compared to trials without saccades (Fig. 2C in velocity, *t*(47) = 3.4, *p* < 0.01; Fig. 2D in magnitude, *t*(47) = 2.55, *p* < 0.05). Saccade and pupil responses were both systematically influenced by stimulation frequency (velocity: Sac: Fig. 2E, R = 0.75, Pupil: Fig. 2F, R = 0.57; magnitude: Sac: Fig. 2I, R = 0.69, Pupil: Fig. 2J, R = 0.61, all *p* < 0.001). Consistent with the common drive hypothesis, trial-by-trial correlational analyses collapsed across all frequency conditions from each stimulation site (N = 24) showed a reliable positive correlation between pupil and saccade responses (histograms of the correlation coefficients are shown in Fig. 2G, K), and the mean correlation coefficient was 0.36 in velocity with significant correlations (*p* < .05) obtained for 58% (14/24) of the stimulation sites (Fig. 2G, parried t-test of *R*-values against zeros: *t*(23) = 7.6, *p* < 0.001), and 0.29 in magnitude with significant correlations obtained for 67% (16/24) of the stimulation sites (Fig. 2K, *t*(23) = 7.1, *p* < 0.001). Again, these results were not an artifact of inaccurate measurement of pupil size because pupil responses relative to the return saccade onset were clearly modulated by stimulation frequency with larger dilation observed for higher stimulation frequencies (Supplementary Figure 2).

**Figure 2.**
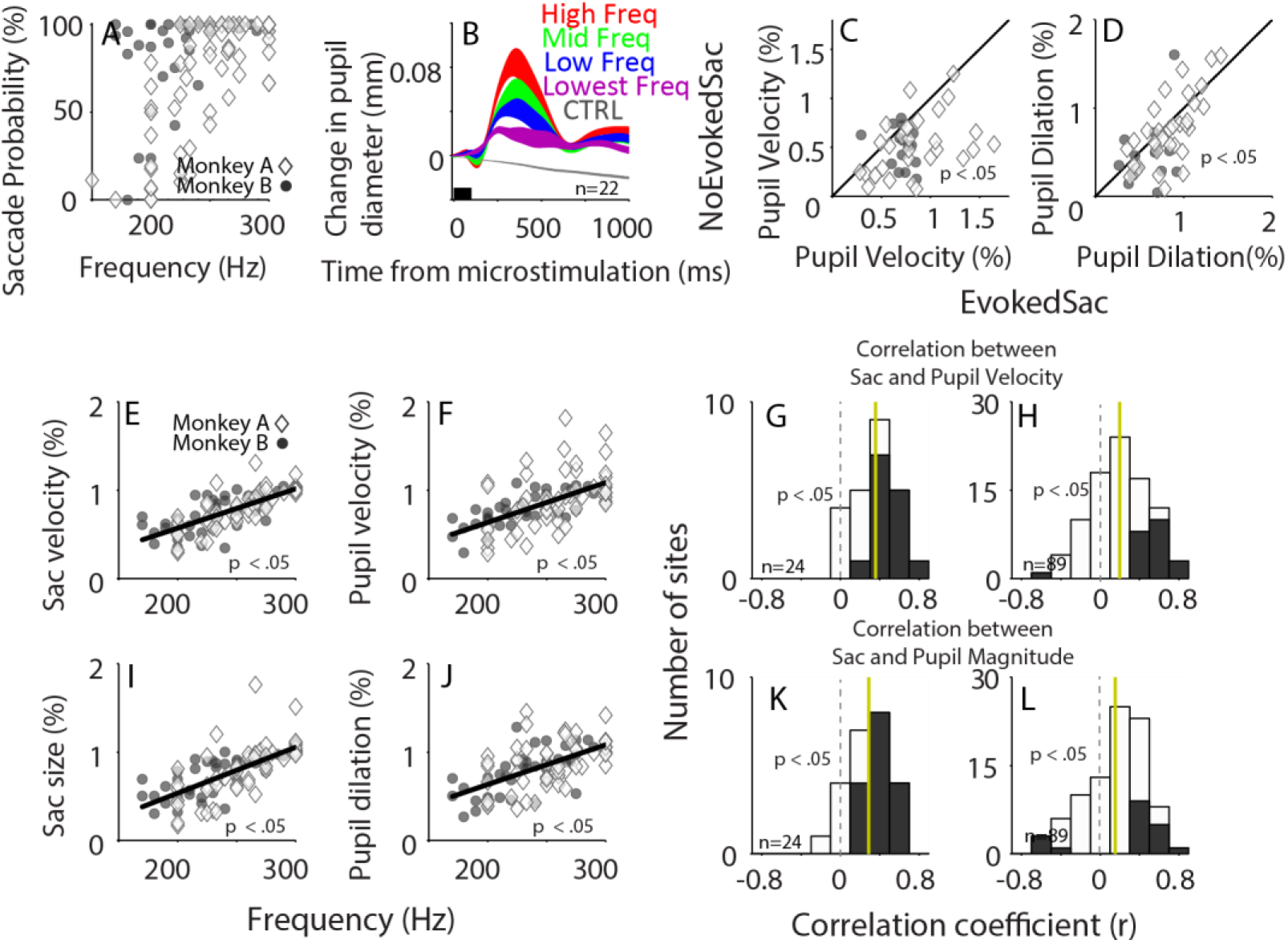
Summary of SCi microstimulation on saccade and pupil modulation. **(A)** Correlation between stimulation frequency and evoked saccade probability. **(B)** Pupil responses after microstimulation in four highest frequency and control (no microstimulation) conditions on data collapsed across monkeys and stimulation sites (n=22). **(C,D)** Pupil responses collapsed across monkeys and stimulation sites (n=24) between trials with evoked saccades and trials without evoked saccades in **(C)** velocity and **(D)** magnitude. **(E,F,I,J)** Effect of stimulation frequency on evoked saccade and pupil responses in **(E,F)**velocity and **(I,J)** magnitude. **(G,K)** Distribution of correlation coefficients for the relationship between evoked saccade and pupil responses in **(G)** velocity and **(K)** magnitude (n = 24). **(H,L)** Distribution of correlation coefficients for the relationship between evoked saccade and pupil responses in each frequency condition in velocity **(H)** and magnitude **(I)** (n = 89). In **A,E,F,I,J**, Black lines indicate the regression line. In **B**, the black bar on X-axis indicates the time line of microstimulation. The shaded colored regions surrounding the pupil response represent ± standard error range for different conditions. In **G,H,K,L**, the vertical black dotted and yellow solid line represent a zero and median value of correlation coefficient. Filled-bars indicate sites with statistically significant correlation (*p* < 0.05). n: number of sites. CTRL: control (no stimulation).

To further examine the correlation between saccade and pupil responses from individual frequency conditions at each site, we split trials according to median saccade peak velocity or magnitude. Although the variation across trials was reduced in a given frequency condition, trials with larger saccade responses still had larger pupil responses (large saccade velocity: 0.85 mm/s pupil velocity; small saccade velocity: 0.73 mm/s pupil velocity; *t*(88) = 4.94, *p* < 0.001; magnitude: large saccade size: 0.095 mm diameter; small saccade size: 0.077 mm diameter; *t*(88) = 5.58, *p* < 0.001). Figure 2H and L show summary histograms of correlation coefficients for all conditions, revealing in general positive correlations between saccade and pupil responses in velocity (significant positive correlations for 24% (21/89) of the individual stimulation/conditions; mean correlation coefficient of 0.2, *t*(88) = 6.5, *p* < 0.001) and in magnitude (significant positive correlations for 17% (15/89) of the stimulation/conditions; mean correlation coefficient of 0.15, *t*(88) = 4.6, *p* < 0.001), though there were a few sites having significant negative correlations (velocity: velocity: 1%, 1/89; magnitude: 4%, 4/89). Together, these results demonstrate positive trial-by-trial correlations between saccade and pupil responses evoked by the SCi microstimulation. Similar results were obtained with varying stimulation duration (Supplementary Figure 3).

### Modulation of stimulation frequency and saccade response on the pupil response

To estimate the contribution of stimulation frequency and saccade responses to pupil dilation, we performed a multiple linear regression analysis of velocity and magnitude across our sample of stimulation sites in two monkeys (data from Fig. 2E,F,I,J) using stimulation frequency (F) and saccade responses (S) as independent variables in the analysis.

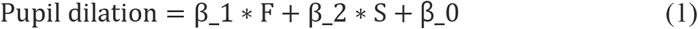

β_0, β_1, β_2 were constant linear weights generated by the model.

The results yielded β_1 = 0.0007, *p* = 0.40; β_2 = 0.8475, *p* = 1.5e-7; R = 0.71 in the velocity analysis, and β_1 = 0.0017, *p* = 0.025; β_2 = 0.5431, *p* = 2.1e-7; R = 0.73 in the magnitude analysis, suggesting that pupil responses were significantly predicted by the saccade response in both velocity and magnitude analyses, but only significantly predicted by stimulation frequency in magnitude analysis, These results imply that the pupil response is more reliably accounted by the saccade response.

### Site-specific effects in saccade and pupil responses

The SCi is organized into a retinotopically coded motor map, and different sites evoke different amplitude and direction saccades ^25^ because different sites project disproportionally to the omnipause and burst neurons ^41–43^. Although unidentified, this pattern of anatomical relationship may also be present in the SCi projections to the pupil control circuit; that is, regions in the SCi map that code for larger amplitude saccades (i.e., caudal SC) may also code for larger pupil dilation, yielding more and/or stronger projections from the caudal SC to the pupil control circuit. Because pupil size is unidimensional, we used eccentricity of the response field determined by the vector of saccade elicited with suprathreshold SCi stimulation (see Materials and Methods) to investigate the relationship between pupil dilation magnitude and evoked saccade size. The magnitude of the evoked pupil dilations was correlated with RF eccentricity: larger dilation and faster velocities were indeed observed for sites with larger vectors of evoked saccades (Fig. 3A in magnitude: R = 0.73, *p* < .001; and 3B in velocity: R = 0.78, *p* < .001). Therefore, our microstimulation results demonstrate the site-specific modulation of both saccade and pupil responses, which is consistent with the common drive hypothesis.

**Figure 3.**
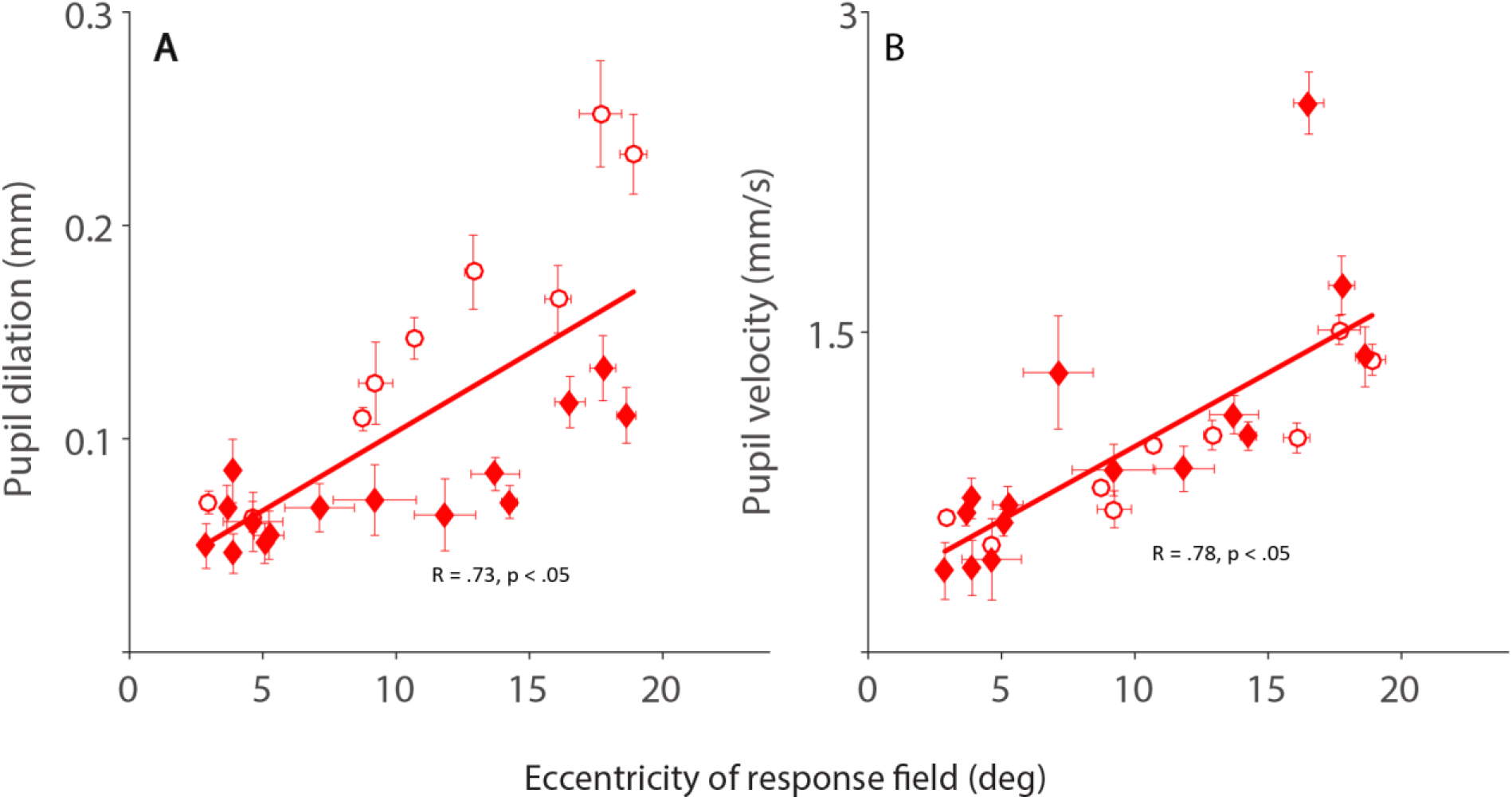
Effects of stimulation-evoked pupillary response as a function of the eccentricity of stimulation site. Correlation between the eccentricity of stimulation site and evoked pupillary response in the three highest frequency conditions in **(A)** magnitude and **(B)** velocity (n = 24). Lines indicate the regression line. Monkey A: ◊, Monkey B: ○.

### Modulation of pupil but not saccade responses by background luminance

The major function of changes in pupil size is to regulate the amount of light projected onto the retina ^29,30^. However, whether global luminance signals play a role in modulating the SC is yet to be explored. Varying background luminance changes the balance of activity in the pupil control pathway ^29^. The pupil responses evoked by the SCi should thus be affected. More importantly, if the SCi is also involved in luminance encoding as suggested by the results of Hannibal et al. ^31^, then varying background luminance should also modulate saccade responses evoked by SCi microstimulation. To investigate the influences of background luminance on pupil and saccade responses evoked by SCi microstimulation, monkeys performed the same fixation task described above, However, this time background luminance was varied, and microstimulation parameters were fixed for each recording site (stimulation duration: 100 ms; stimulation frequency range: 240 – 300 Hz). Changes in background luminance systematically changed pupil diameter, as shown in Fig. 4 for an example stimulation site. Consistent with the literature ^29^, increasing background luminance systematically reduced baseline pupil diameter with mean value of 3.07, 2.47, 2.2, 1.79, 1.56 mm in the 2.5, 5, 10, 20, 40 cd/m^2^ background conditions, respectively (Fig. 4A; differences were all statistically significant between any of two conditions according to bootstrapping, see Materials and Methods). Importantly, pupil responses evoked by SCi microstimulation were also systematically modulated by background luminance levels (Fig. 4B). Increasing background luminance produced lower pupil velocities (Fig. 4C, low to high background luminance: mean value: 1.42, 1.26, 1.07, 0.66, 0.85 mm/s) and less dilation (Fig. 4D, low to high background luminance: mean value: 0.38, 0.35, 0.25, 0.16, 0.16 mm). In contrast, saccade responses were not systematically modulated by background luminance, with mean peak velocities 583, 572, 581, 498, 546 deg/s (Fig. 4E) and saccade sizes 13.3, 13.6, 14.3, 12.3, and 13.6 deg (Fig. 4F) in the 2.5, 5, 10, 20, 40 cd/m^2^ background conditions, respectively, although some comparisons were statistically significant.

**Figure 4.**
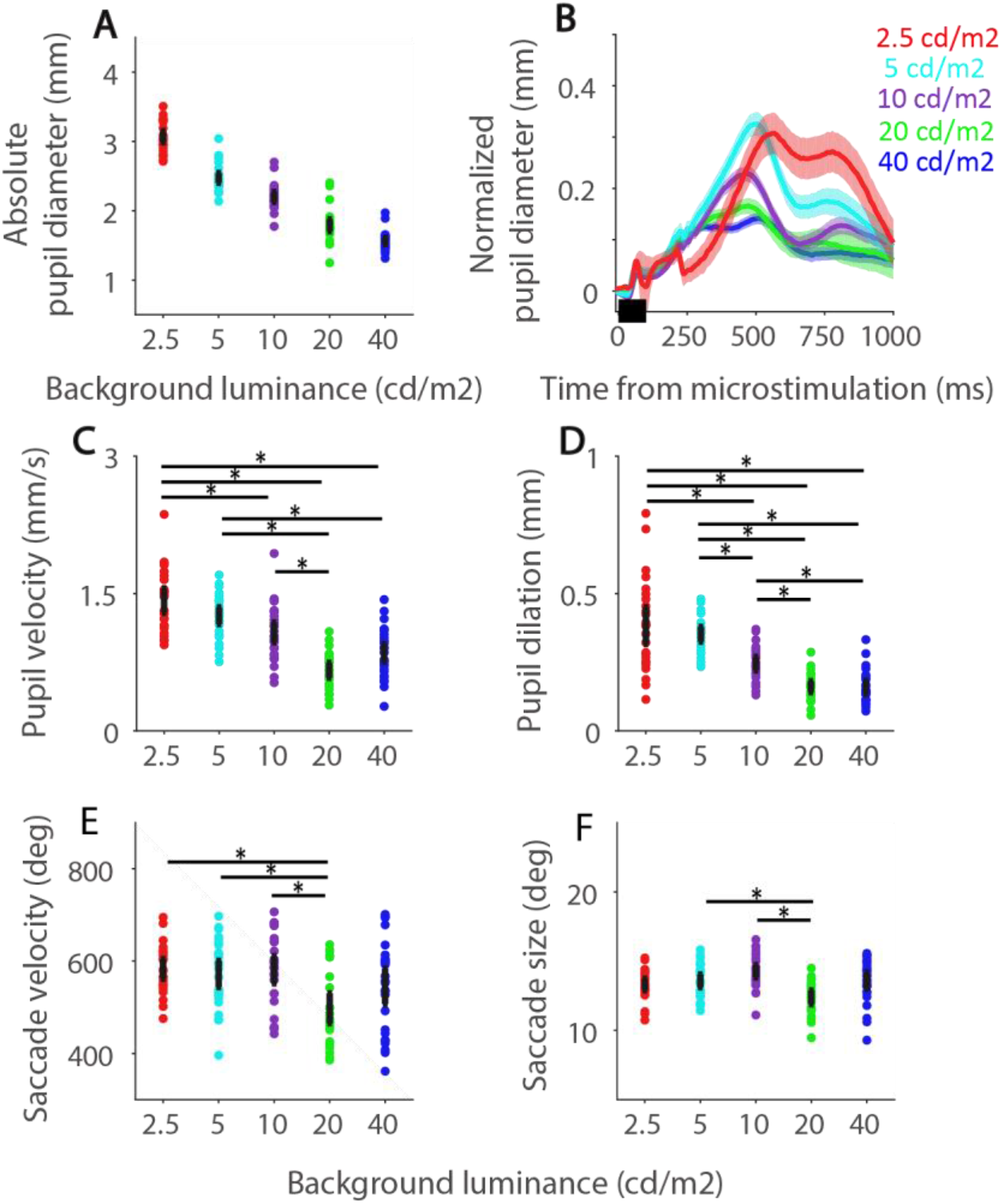
Examples showing background luminance effects on saccade and pupil responses. **(A)** Effects of background luminance on absolute pupil diameter at an example site. **(B)** Pupil responses after microstimulation in five background luminance conditions. **(C-D)** Pupil responses after microstimulation in different background luminance conditions with respect to **(C)** velocity and **(D)** magnitude. **(E-F)** Saccade responses after microstimulation in different background luminance conditions with respect to **(E)** velocity and **(F)** magnitude. In **A,C-F**, the thick black and gray circles represent an average and individual trial values of measurements. In **B**, the black bar on X-axis indicates the time line of microstimulation. The shaded colored regions surrounding the pupil response or error-bar represent ± 95% confidence interval for different conditions derived from the bootstrap analysis. n: number of sites. *: differences were statistically significant according to bootstrapping.

Figure 5 summarizes the effects of SCi microstimulation on saccade and pupil responses in different background conditions (2.5, 20, 40 cd/m^2^) collapsed across all stimulation sites tested with luminance changes from 2 monkeys (N = 16) (note that this was minimal data set for all possible sites and some sites, e.g. Fig 4, had more than 3 luminance values). Pupil diameter (Fig. 5A, sac: F(2,30) = 1417.86, *p* < 0.001; *Bonferroni-corrected comparisons:* all comparisons, *p* < 0.05) prior to microstimulation was clearly modulated by background luminance, with absolute pupil sizes 2.45, 1.35, and 1.13 mm for 2.5, 20, 40 cd/m^2^, respectively. Figure 5B shows the normalized pupil diameter produced by SCi microstimulation for three background conditions (stimulation versus no-stimulation), revealing larger responses for lower luminance conditions. To quantify these orienting responses, we normalized these responses (see Materials and Methods). As predicted, evoked pupil responses were systematically modulated by background luminance (Fig. 5C: velocity: F(2,30) = 44.163, *p* < 0.001, all comparisons *p* < 0.01; Fig. 5D: magnitude, F(2,30) = 221.934, *p* < 0.001, comparisons between 2.5 cd/m^2^ and others, *p* < 0.05), with larger pupil dilation evoked with lower background luminance. In contrast, the probability of evoked saccades was not influenced by different background luminance (2,5, 20, 40 cd/m^2^: 94, 92, 91 %, F(2,30) = 0.538, *p* = 0.59). Moreover, neither the velocity (Fig. 5E: F(2,30) = 1.486, *p* = 0.24, all comparisons, *p* > 0.37) nor magnitude (Fig. 5F: F(2,30) = 0.999, *p* = 0.38, all comparisons, *p* > 0.5) of the saccades were modulated by background luminance levels. These results characterize the influence of luminance signals on pupil, but not, saccade responses evoked by SCi microstimulation.

**Figure 5.**
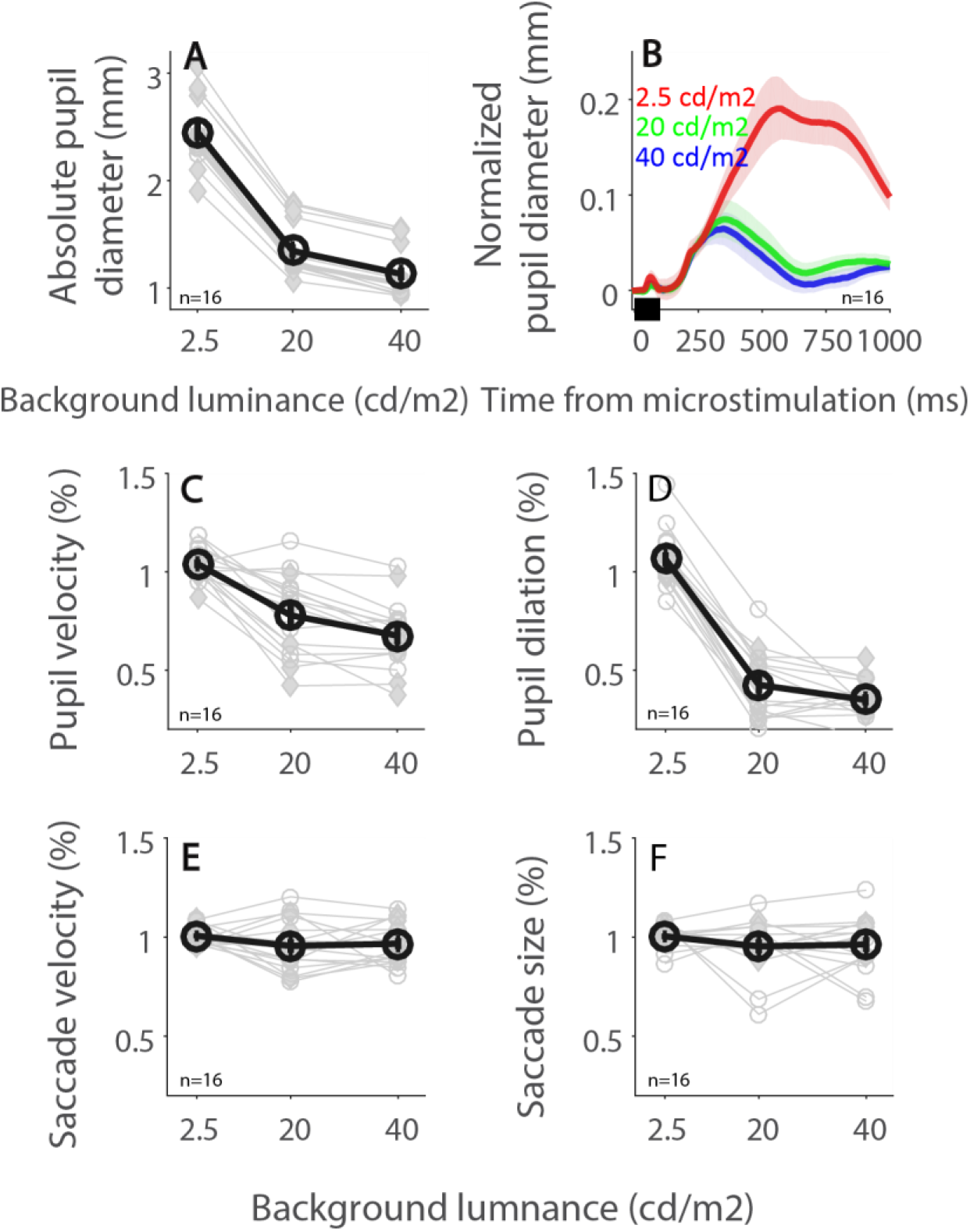
Summary of background luminance on saccade and pupil responses. **(A)** Effects of background luminance on absolute pupil diameter collapsed across monkeys and stimulation sites (n=16). **(B)** Pupil responses after microstimulation for three background luminance conditions. **(C-D)** Pupil responses after microstimulation for different background luminance conditions in **(C)** velocity and **(D)** magnitude. **(E-F)** Saccade responses after microstimulation in different background luminance conditions in **(E)** velocity and **(F)** magnitude. In **A,C-F**, the thick black and gray circles represent an average and individual site values of measurements. In **B**, the black bar on X-axis indicates the time line of microstimulation. The shaded colored regions surrounding the pupil response represent ± standard error range for different conditions. n: number of sites. Monkey A: ◊, Monkey B: ○.

## Discussion

The goal of this study was to understand the role of the SCi in the coordination between saccades and pupil dilation during the orienting response, and to determine whether background luminance interacts with these evoked responses (Fig. 1A). Varying stimulation frequency systematically modulated the evoked saccade as well as the pupil responses, with larger saccade and pupil responses observed for higher stimulation frequencies. Trial-by-trial correlation was also observed: trials with larger saccade responses had larger pupil responses. Furthermore, microstimulation of SCi sites, which produced larger amplitude saccades, also evoked larger pupil dilation. These results together demonstrate saccade and pupil movements are coordinated through the SCi, likely via a shared common output signal derived from the SCi. However, varying background luminance only influenced evoked pupil responses, implying that luminance information was not processed directly in the SCi, although the SC does receive direct retinal projections ^31^.

### Circuits for saccade and pupil control by the superior colliculus

The SCi integrates sensory, motor, and cognitive signals from various cortical and subcortical areas and projects directly to the premotor brainstem circuitry to initiate the orienting response ^10,12^. Although the role of the SCi in the control of saccade and pupil responses has been developed, it has yet to be determined whether the same SCi efferent projection drives both saccade and pupil responses (the common drive hypothesis), or alternatively, that saccade and pupil responses are mediated by different sets of axons emerging from the SCi. The connections between neurons in the SCi and brainstem reticular formation for control of saccades is well-established ^44–46^. The central mesencephalic reticular formation (cMRF) is one of the collicular targets involved in horizontal saccade control ^47^, and research using intra-axonal recording and horseradish peroxidase injection has revealed that saccade-related long-lead burst neurons in the SCi with axon collaterals terminating in the cMRF also commonly project to the pontine premotor centers for saccade control ^48,49^. Moreover, it has recently shown that cMRF also projects directly to the Edinger–Westphal nucleus ^50^, a critical structure in the pupil control circuit ^10,12^. In addition, the superior colliculus projects directly, and indirectly via the cMRF, to the medullary reticular formation ^51^, and so may influence preganglionic sympathetic motoneurons via this route. However, connections to the motoneurons controlling dilation via these routes remain to be demonstrated. These results and trial-by-trial correlations together suggest that the same efferent projection derived from the SCi could possibly drive both saccade and pupil premotor circuits to produce coordinated saccade and pupil responses. We found that microstimulation of the SCi produced trial-by-trial correlations between saccade and pupil responses, suggesting that common output signals derived from the SCi drive both down-stream premotor circuits. Although trial-by-trial correlation between saccadic and pupil responses could theoretically be explained by assuming different SCi efferents are activated via current spreading, current spread, anatomical data on SCi efferents supports the common drive hypothesis.

Monkeys returned their eyes back to the central FP (return saccade) after a saccade evoked by SCi microstimulation. The properties of the return saccade are very similar to the evoked saccade, albeit traveling in the opposite direction. Moreover, pupil size should also be modulated in a manner that correlates with the return saccade. Due to the speed of these two saccades, it appears that the pupil effects were combined. Certainly, robust trial-by-trial correlations were still observed even though we only compared dilation to the properties of the stimulus evoked saccades. Future studies are thus needed to explore how a sequence of saccades might modulate pupil response dyanamics.

Different regions in the SC motor map project disproportionally to omnipause neurons and excitatory/inhibitory burst neurons to trigger saccades with different amplitudes ^41–43,52^. Although unidentified, this pattern of anatomical relationship may also be present in the SCi projections to the pupil control circuit; that is, regions in the SCi map that code for larger amplitude saccades (i.e., caudate SC) also code larger pupil dilation, yielding more and/or stronger projections in the caudal SC to the pupil control circuit. Our results support this idea: SC sites that evoked larger size saccades also induced larger pupil dilation. Future research is required to explore the anatomical relationship between different regions in the SC and its connections to the pupil control circuit.

### Role of other brain areas in pupil control

Pupil size is also modulated by other structures in the brain that possibly contribute to correlations observed here between saccade and pupil responses ^53,54^. Mounting evidance has demonstrated that pupil size covaries with neural activity of the locus coeruleus (LC) in behaving monkeys ^20,22,55^, and that LC neurons also discharge phasically to task-relevant sensory stimuli ^56–61^. Furthermore, the cholinergic system also correlates with pupil responses ^21^, and could also modulate saccade responses through changing cholinergic activity within the SC via input from the pedunculopontine tegmental nucleus (PPTN) ^62–64^. Pupil dilation is also evoked by subthreshold microstimulation of the frontal eye fields (FEF) ^19,65^, a structure involved in attention and gaze shifts ^66^, although these studies focused mainly on pupil dynamics, instead of the coordination between saccade and pupil responses. Therefore, it is possible that abovementioned structures are also involved in observed correlations between pupil and saccade responses. Furthermore, the SCi receives inputs from the LC, FEF as well as cholinergic projections from the PPTN ^15–18^. Therefore, it is likely that the signals from LC, FEF, and PPTN are integrated in the SCi to drive coordinated saccade and pupil responses. This idea can also explain why pupil responses are more reliably modulated by saccade responses than by stimulation frequency because the saccade response is the summation of SC inputs (e.g., LC, FEF, cholinergic) and stimulation frequency.

### Global luminance effects on evoked saccades and pupil dilation and functional role of its coordination

The SC receives retinal projections, including ipRGCs projections to both the SCs and SCi ^31^, which are particularly important to luminance encoding and the pupillary light reflex ^32–35^. However, it is unclear whether these luminance signals serve a role in signal processing in the SCi. Varying background luminance changes pupil size by regulating the activity balance between the sympathetic and parasympathetic pathways, with greater activation in the sympathetic system, but smaller activation in the parasympathetic system, at lower levels of luminance ^29,67^. Here, we found that lowering background luminance systematically increased the degree of pupil dilation evoked by SCi microstimulation, but it did not change the evoked saccade responses. This suggests an independent relationship between the SCi and luminance signal processing. Notably, although the global luminance did not modulate saccades evoked by microstimulation, it may influence saccades that are heavily associated with visual processes such as visually-guided saccades, so, future work is required to examine this question.

Pupil dilation can be mediated through inhibiting the parasympathetic system ^68^ and/or activating the sympathetic system ^29,67^. According to the modulation of the balanced activity between the sympathetic and parasympathetic pathways by background luminance, pupil dilation should be mediated mainly by the excitation of the sympathetic pathway under lower luminance conditions, but by the inhibition of the parasympathetic pathway under higher luminance conditions. Because dilation here was greater in lower global luminance, such dilation was likely due to sympathetically-driven dilator pupillae contraction which is reduced by antagonism from increased sphincter pupillae contraction at higher luminance levels.

Larger pupil dilation observed in lower luminance conditions also leads to an intriguing question because pupil dilation should be larger under higher luminance conditions according to the mechanical limit of pupil size. The fact that larger pupil dilations were produced by SCi activation when luminance was lower is intriguing because pupil dilation should theoretically be larger under low light conditions where the pupil is near the mechanical limit of pupil size. Our finding the opposite effect may reflect the functional role of orienting pupil dilation. It has been argured that pupil dilation evoked by salient stimuli serves to slightly increase visual sensitivity ^69^, so the size of evoked pupil dilation should be larger under lower luminance conditions because it is particularly needed to increase visual sensitivity under these conditions. Consistently, the observed pupil dilation produced by SCi stimulation was larger under lower background luminance conditions, which is consistent with our previous results with subthreshold microstimulation ^4^. More interestingly, saccade and pupil responses evoked by the SCi, although different in time course, are coordinated. Does the coordination between saccade and pupil responses serve any functional role for visual processing? Although the current study cannot address this question, a potential benefit may be having a larger pupil size after orientation, i.e., saccades, resulting in efficient processing of a selected target at the beginning of the foveation. Future research is needed to directly examine the functional role of pupil dilation evoked by salient stimuli and mediated by the SC for visual processing.

### Conclusion

The SCi, a hub of sensory and motor processing, integrates sensory and cognitive signals to coordinate the orienting response that includes eye movements and pupil size changes ^10,12^. Here, we demonstrated saccades and pupil dilation evoked by SCi microstimulation were highly correlated, and SCi site-specific effects were observed in both saccade and pupil responses, together supporting the common drive hypothesis, and arguing that the same efferent projection from the SCi drives both saccade and pupil responses. Moreover, varying background luminance only modulated evoked pupil, but not saccade, responses, implying the functional independence of the SCi and luminance signals. Orienting responses are thought to work together to optimize the body for whatever action is required ^69^. Investigation of the various orienting components simultaneously is thus necessary to understand how they are coordinated to optimize performance.

## Materials and Methods

### Animal experimental setup

Experiments were performed on two male rhesus monkeys (*Macaca mulatta*; 10 and 11 kg). The protocols used in this study were approved by Queen’s University Animal Care Committee in accordance with the Canadian Council on Animal Care policies for the use of laboratory animals. The methods of surgical procedures, techniques for extracellular neuronal recording, and data collection have been described in detail previously ^70^. Eye position and pupil size were measured by a video-based eye tracker (Eyelink-1000, SR Research, Osgoode, ON, Canada) at a rate of 1000 Hz with monocular recording (right pupil). Pupil area values recorded from the eye tracker were converted to pupil diameter (see details in ^5^. Stimulus presentation and data acquisition were controlled by a UNIX based real-time data control system (REX) (Hays et al., 1982). Spikes, eye position, and pupil diameter were recorded in a multichannel data acquisition system (Plexon). Stimuli were presented on a CRT monitor at a screen resolution of 1024 × 768 pixels (75Hz non-interlaced), subtending a viewing angle of 54 × 44 deg.

### Procedure, SC recording, and stimulation

Monkeys were seated in a primate chair with their heads restrained facing the video monitor. We lowered tungsten microelectrodes (impedance: 0.1–1 MΩ, Frederick Haer) to locate the SC. We mapped the visual response fields of the SC using a visual mapping task ^36^, and identified the depth of the SCi (intermediate layers of the superior colliculus) using a delayed saccade task. Once the SCi was localized, it was microstimulated (250-300 Hz pulse train for 100 ms with alternating 0.3 ms anode plus 0.3 ms cathode pulses), and threshold for saccades was determined when the stimulation current in the SCi evoked saccades 50% of the time (range: 5-50 µA). The optimal locations of the response fields of SCi neurons were in close agreement with the vector of eye movement elicited with suprathreshold SCi stimulation, and ranged between 3° and 20° eccentricity.

### Experimental task

The monkeys were trained to perform a simple fixation task. They had to maintain gaze within 1.5° of a central fixation point (FP, 0.5° diameter; 20 cd/m^2^, isoluminant color of the background) at the center of the screen on a gray background (20 cd/m^2^) for a few seconds to obtain a liquid reward. After the monkey maintained fixation for 1–1.5 s, a train of stimulation pulses was delivered on 50% of the trials, and the monkeys had to maintain fixation for another 1.5–2 s, regardless of microstimulation. Saccades were often evoked after electrical stimulation (latencies usually less than 100 ms). Monkeys had to move their eyes back to the FP within 500 ms after microstimulation, and both monkeys usually moved their eyes back to the FP within 300 ms after microstimulation.

Two microstimulation experiments were conducted. In the first experiment, stimulation frequency was manipulated (100 ms stimulation train duration). After determining the saccade threshold current, usually at 300 Hz stimulation, we systematically varied the frequency of stimulation, ranging from 150 to 300 Hz, and used ~150 % of the saccade threshold current to regularly evoke saccades, particularly under high frequency stimulation. At each stimulation site, we microstimulated at 3-8 different frequency levels. The order of frequency levels across blocks was varied across days. Microstimulation was delivered to 24 sites (9 and 15 in monkeys A and B, respectively).

In the second experiment, the level of global luminance was manipulated by changing the background luminance level of the computer screen. We thus refer to background lumiance of the screen as global luminance. Stimulation levels were fixed across various background luminance conditions (> 250 Hz stimulation frequency and 100 ms stimulation duration). The monkey had to maintain gaze within 1.5° of a central FP (0.5° diameter; isoluminant color difference from background) at the center of the screen with background luminance of 2.5, 20, or 40 cd/m^2^. The order of luminance levels across blocks was varied across days. Microstimulation was delivered to 16 sites (6 and 10 in monkeys A and B, respectively) at 3-5 background levels (2.5, 5, 10, 20, 40 cd/m^2^). There were at least 20 correct trials in all conditions.

### Data analysis

Trials with blinks or an eye position deviation of more than 1.5° from the central FP or with the detected saccades (> 2°) during the required period of central fixation were excluded from analysis except for microstimulation-related (evoked and return) saccades. The recorded pupil size depends on the subject's gaze angle in a video-based eye-tracker so that pupil-size data can be distorted by eye movements and eccentric eye positions. Because of applying suprathreshold microstimulation, saccades were often evoked after stimulation and eye position returned to center within 300 ms. To maintain an accurate measure of the pupil, we mainly used pupil peak dilation and velocity measures because they regularly occurred after 300 ms of microstimulation, and at this time the eyes had returned to the fixation point, so that the pupil could be accurately measured. The difference in the efferent delays of the saccade and pupillary responses are primarily caused by the different muscle types being innervated: smooth muscle controlling the pupil and extraocular fast twitch skeletal fibers for the saccades. Nevertheless, the pupil response is temporally dissociated from the saccade response (see Fig. 1B for an example).

Because the pupil response is consensual ^4,5^, only pupil diameter of the right eye was recorded for data analysis. Following the procedures of baseline-correction used previously ^4,6^, original pupil diameter values were subtracted from the baseline pupil diameter value determined by averaging pupil size from 200 ms before to the onset of microstimulation for each trial. To normalize pupil and saccade measurements across conditions for population analyses (frequency, background luminance), all measured values were divided by the median value of the highest frequency condition (lowest luminance level for the other experiments). To be included for condition-based analyses, each condition (frequency/background luminance) had to have more than 15 remaining trials, except where indicated. Evoked saccade reaction time (SRT) was defined as the time after microstimulation to the first saccade away from central fixation that exceeded 30°/s. The highest three frequency conditions were selected for some analyses to examine the relationship between saccade and pupil responses because there were more numerous trials with evoked saccades in the higher frequency conditions.

Results are shown as mean ± SEM. We performed correlational analyses and a two-tailed student *t* test except where indicated. Bayesian *t* tests, where appropriate, were also performed to inform statistical significance for pairwise comparisons, with a scale factor r = 0.707 ^71^. Moreover, Cohen’s *d*, where appropriate, was calculated to estimate effect size ^72^. In the background luminance experiment, we used a bootstrap method to inform the statistical significance of the comparison by performing a random sampling of pupil values derived from each recording trial with 1000 repetitions ^6^. This resulted in a normally distributed cluster of points centered on the mean of selected pupil values (clusters not shown, normal distribution was verified by the Kolmogorov-Smirnov test). Statistical tests were performed using JASP Team (2019) and MATLAB (The MathWorks Inc., Natrick, MA, USA).

## Acknowledgement

This work was supported by Canadian Institutes of Health Research Grant (MOP-FDN-148418) and the Canada Research Chair Program to DPM, and grants from Taiwan Ministry of Science and Technology (108-2410-H-038-002-MY3; 109-2636-H-038-005) to CW. We thank Ann Lablans, Brittney Armitage-Brown, and Mike Lewis for outstanding technical assistance.

